# CRISPR-Targeted CAR Gene Insertion Using Cas9/RNP and AAV6 Enhances Anti-AML Activity of Primary NK Cells

**DOI:** 10.1101/2021.03.17.435886

**Authors:** Meisam Naeimi Kararoudi, Shibi Likhite, Ezgi Elmas, Kenta Yamamoto, Maura Schwartz, Kinnari Sorathia, Marcelo de Souza Fernandes Pereira, Raymond D Devin, Justin M Lyberger, Gregory K Behbehani, Nitin Chakravarti, Branden S. Moriarity, Kathrin Meyer, Dean A. Lee

## Abstract

Human peripheral blood natural killer (NK) cells have intense antitumor activity and have been used successfully in several clinical trials. Modifying NK cells with a chimeric antigen receptor (CAR) can improve their targeting and increase specificity. Recently, we described an efficient method for gene targeting in NK cells using Cas9/ribonucleoprotein (RNP) complexes. Here we combined this approach with single-stranded (ss) or self-complementary (sc) Adeno-associated virus (AAV)-mediated gene delivery for gene insertion into a safe-harbor locus using a wide variety of homology arms for homology repair (HR) and non-homologous directed CRISPR-assisted insertion tagging (CRISPaint) approaches. For proof-of-concept, we generated mCherry-expressing primary NK cells and determined that sc vectors with 300bp homology arms had optimal transduction efficiency. Then, we generated CD33-targeting CAR NK cells with differing transmembrane and signaling domains (CD4/4-1BB+CD3ζ and NKG2D/2B4+CD3ζ) and expanded them on CSTX002 feeder cells. Expansion kinetics were unaltered and the expanded NK cells maintained high CAR expression (mean 68% CAR+). The CD33-CAR-NK cells showed increased activation markers and enhanced antileukemic activity with improved killing kinetics against CD33-positive acute myeloid leukemia (AML) cell lines and primary samples. Using targeted sequencing we demonstrated the accuracy of CAR gene insertion in human primary NK cells genome. Site-directed insertion using RNP and scAAV6 is an efficient method for stable genetic transfer into primary NK cells that has broad potential for fundamental discovery and therapeutic applications.

## Introduction

Human primary NK cells have been tested in numerous clinical trials demonstrating a high safety profile and evidence of clinical benefit for patients with cancer, which has been most widely applied to acute myelogenous leukemia (AML). Enhanced targeting of NK cells to AML through CD33 targeting has been shown with antibody Fc modifications^1, 2^ or fusion to alternative activation domains as bi- or tri-specific NK cell engagers^3, 4^. CD33-targeting by T cells has been enabled by CD33-targeting chimeric antigen receptors (CAR)^5–7^, but clinical application has been hindered by concerns of long-term suppression of hematopoiesis with CAR-T persistence.

Gene modification of NK cells to enable stable expression of a CAR can also improve their antitumor activity^8^. However, gene modification of human peripheral-blood derived NK cells (PB-NK) using viral or non-viral vectors has been challenging due to robust foreign DNA- and RNA-sensing mechanisms that limits the efficiency of these gene delivery methods^8^. To overcome this limitation, mRNA-based gene delivery has been tested in PB-NK cells, but only allows for transient expression of transgenes^9^. We recently demonstrated highly efficient gene knockout in human primary NK cells by electroporating Cas9/ribonucleoprotein complexes (Cas9/RNP)^10–12^. After Cas9 introduces a DSB, two independent and innate DNA repair mechanisms may be employed to repair the break: homologous recombination (HR) or non-homologous end-joining (NHEJ). In the presence of a DNA template encoding a gene of interest, the exogenous gene can be integrated into the Cas9-targeting site using either of these repair mechanisms^13^. There are several ways to provide the DNA template, including viral and non-viral methods. In non-viral approaches, the single-stranded or double-stranded DNA template is typically electroporated along with Cas9/RNP^14^, however, it typically has a lower efficiency in comparison to viral transduction. For viral gene delivery, adeno-associated viruses (AAV), including AAV6, have been used safely as delivery vectors in clinical trials for primary immune cells, including T-cells ^15, 16^.

The Cas9/RNP-AAV approach has not been described in human primary NK cells. Here, we set to optimize this approach wherein Cas9/RNP electroporation in primary human NK cells is followed by a DNA template encoding a transgene, with or without homology arms for Cas9 targeting site, delivered using single stranded or self-complementary AAV6. Using this approach, we achieved simple yet highly efficient generation of stable transgene-modified human primary NK cells, including two CAR-NK cells which showed enhanced anti-AML activity. The gene modified NK cells generated by the AAV-Cas9/RNP platform has utility for clinical applications such as CAR expression for antigen specific cancer immunotherapy, and for studying NK cell biology.

## Results

### Expansion of NK cells provides optimal conditions for gene insertion

The DNA modifying and repairing enzymes required for NHEJ and HR are different. NHEJ, which is essential for CRISPR-assisted insertion tagging (CRISPaint), is a LIG4-dependent process, while BRCA1 and BRCA2 are essential for HR^31, 32^. We have shown that expansion of NK cells on feeder cells expressing membrane-bound IL-21 (FC21) induces broad changes in gene expression^18, 19^. Therefore, we analyzed the expression level of genes essential for HR and NHEJ in freshly isolated NK cells and after seven days of stimulation (Day 7 expanded cells) to determine the timepoint at which these mechanisms would be most active. RNA-seq showed that the expanded NK cells had highly increased expression of BRCA1 and BRCA2 in comparison to naïve NK cells, and there was no significant change in LIG4 expression (Figures 1a and 1b), suggesting optimal conditions for either HR or NHEJ-directed gene insertion in expanded NK cells.

**Figure 1:**
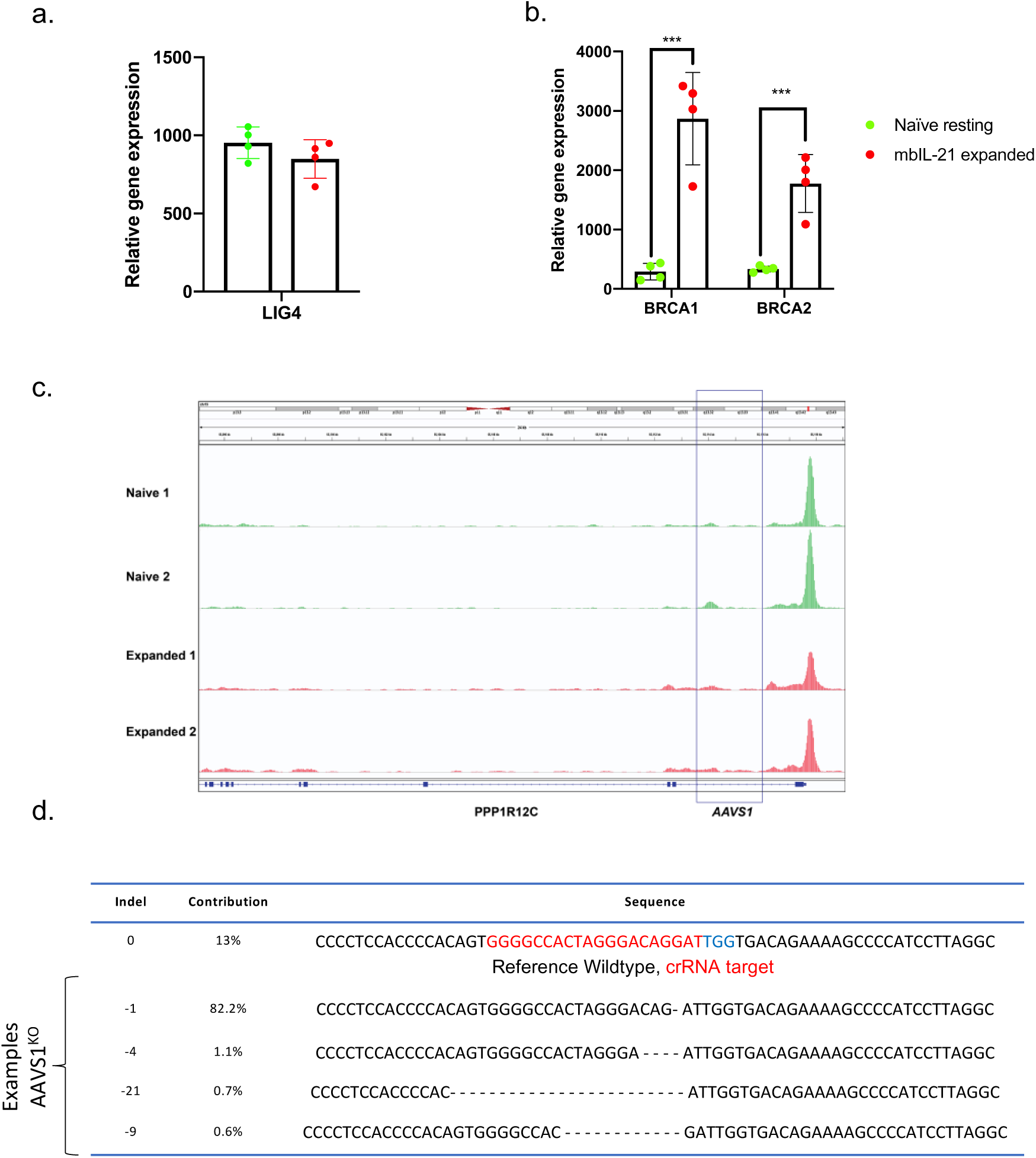
Efficient CRISPR targeting of AAVS1 in mbIL-21 expanded human primary NK cells. Relative gene expression level of NHEJ-related genes **(a)** and HR-related genes **(b)** in naïve and mbIL-21 expanded NK cells, ***P < 0.001 for all comparisons (n=4). **c,** ATAC-seq data shows that AAVS1 has a similar chromatin accessibility between freshly isolated (Naïve), mbIL-21 expanded NK cells (n=2). **d.** Efficiency of Cas9/RNP-mediated targeting of AAVS1 in NK cells. Data are shown as mean ± s.d. P values are from 2way ANOVA.

### Targeting a genomic safe harbor for gene insertion

For gene insertion in NK cells, we chose the *adeno-associated virus integration site 1 (AAVS1),* which is an exemplary genomic safe harbor locus within the *phosphatase 1 regulatory subunit 12C (PPP1R12C)* gene^13, 33^. Chromatin accessibility of AAVS1 was similar in naïve and Day 7 expanded NK cells (n=2) as determined by ATAC-seq (Figure 1c). Next, AAVS1 was targeted by electroporation of Cas9/RNP into Day 7 expanded NK cells (Supplementary Figure 1a) ^11^. After 48 hours, the frequency of insertions and deletions (Indels) in CRISPR-edited NK cells was determined using Inference of CRISPR Edits (ICE) using primers flanking the AAVS1 locus (Supplementary Table 1) ^22^. The ICE results showed that up to 85% of CRISPR modified NK cells had at least one indel at the AAVS1 Cas9-targeting site (Figure 1d). Using the Calcein-AM assay^11^, we observed no difference between wild type and CRISPR-modified AAVS1^KO^ NK cells in their killing ability against Kasumi-1, an acute myeloid leukemia (AML) cancer cell line, suggesting that genome modifications at this locus do not interfere with the ability of NK cells to target cancer cells (Supplementary Figure 1b).

### Gene insertion in primary human NK cells using single-stranded AAV6 and Cas9/RNP

To compare the efficiency of gene insertion across DNA-repair mechanisms, we generated a parallel series of AAV6 vectors suitable for HDR-mediated gene insertion (Figure 2a) using both single-stranded and self-complementary designs with varying homology arm lengths, and for NHEJ-mediated gene insertion with CRISPaint containing PAMgPAMg sequences (Figure 2b). To maximize HDR-mediated gene insertion, we identified homology arms (HA) for the right and left sides of the flanking regions of the Cas9 targeting site in the AAVS1 locus, cloned these together with the mCherry gene into the backbone of a single-stranded AAV plasmid, and packaged this construct into the AAV6 viral capsid (Figure 2a) ^16, 34, 35^. It has been shown that the efficiency of recombination increases as the length of HAs increases^36–40^. Therefore, for the ssAAV backbone, we used the longest possible length of the left and right HA for mCherry (800bp-1000bp of HAs). The constructs also contained a splice acceptor downstream of the transgene to improve the transcription of the mCherry gene (Figure 2a). Electroporation of the NK cells with Cas9/RNP targeting AAVS1 followed 30 minutes later by AAV transduction (Figure 2c)^10^ resulted in 17% (300K MOI) and 19% (500K MOI) mCherry positive NK cells (Figure 3a). We further expanded these cells for one week using FC21, enriched the mCherry positive cells by FACS sorting, and did not see any reduction in the expression level of mCherry during an additional 30 days of expansion (Figures 3a, 3b, and 3c) demonstrating stable integration.

**Figure 2:**
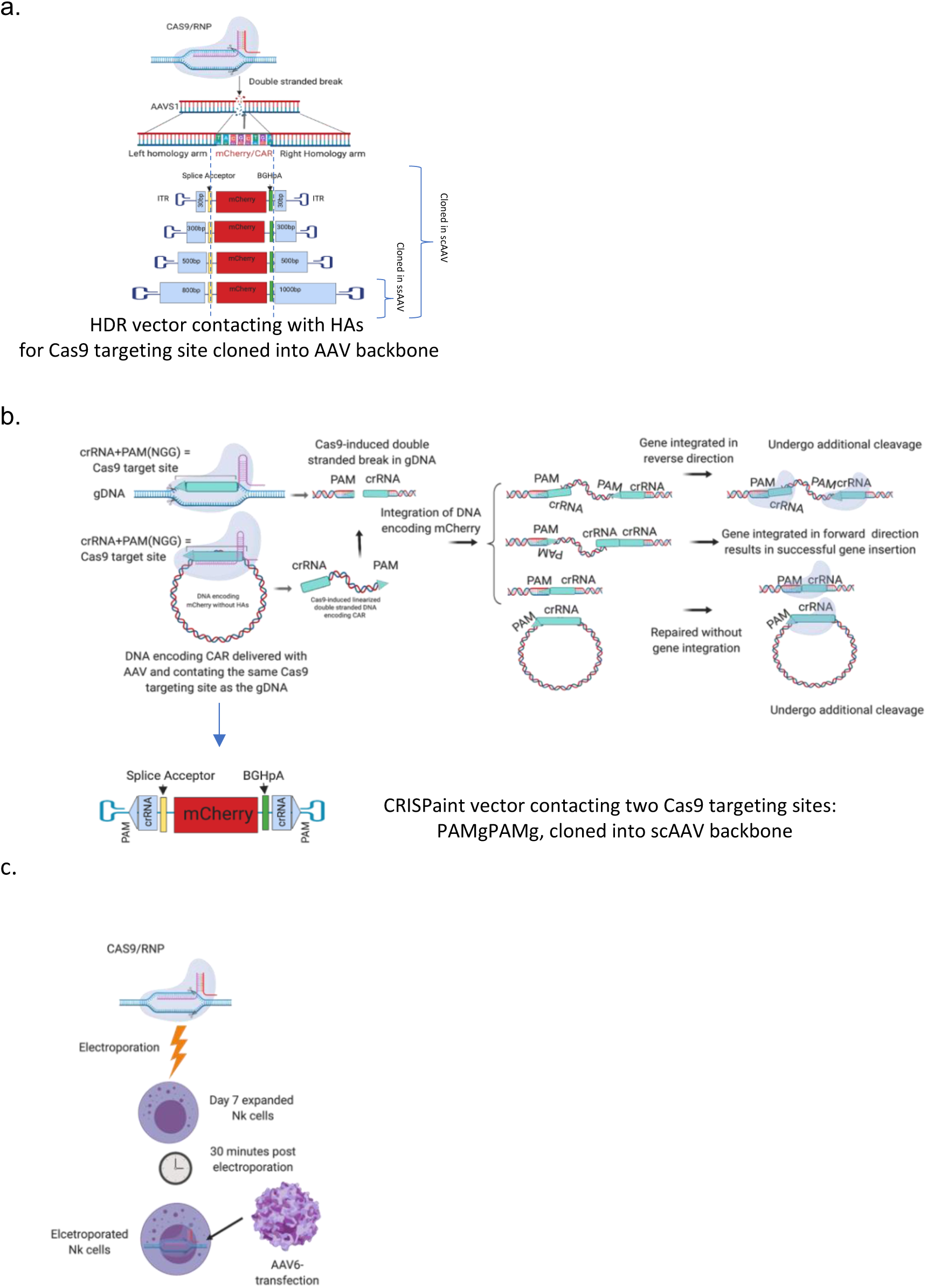
Constructs of mCherry encoding DNA for insertion into AAVS1 through HR and CRISPaint. **a**, Cas9/RNP introduces DSB in AAVS1, DNA encoding gene of interest can be integrated into NK cells through HR with optimal length of HAs. The schematics show the constructs design for integration of DNA encoding mCherry with HAs between 30-1000bp for Cas9 targeting site in AAVS1 and cloned in ssAAV6 and/or scAAV6 backbone. **b,** Top, schematics of how CRISPaint gene insertion works through homology independent DNA repair pathway. Bottom, schematic of construct design for insertion of DNA encoding mCherry through CRISPaint and cloned in scAAV. **c,** Schematics of workflow to electroporate Cas9/RNP and transduce day seven mbIL21 expanded IL2-stimulated NK transduced with AAV6 for gene delivery.

**Figure 3.**
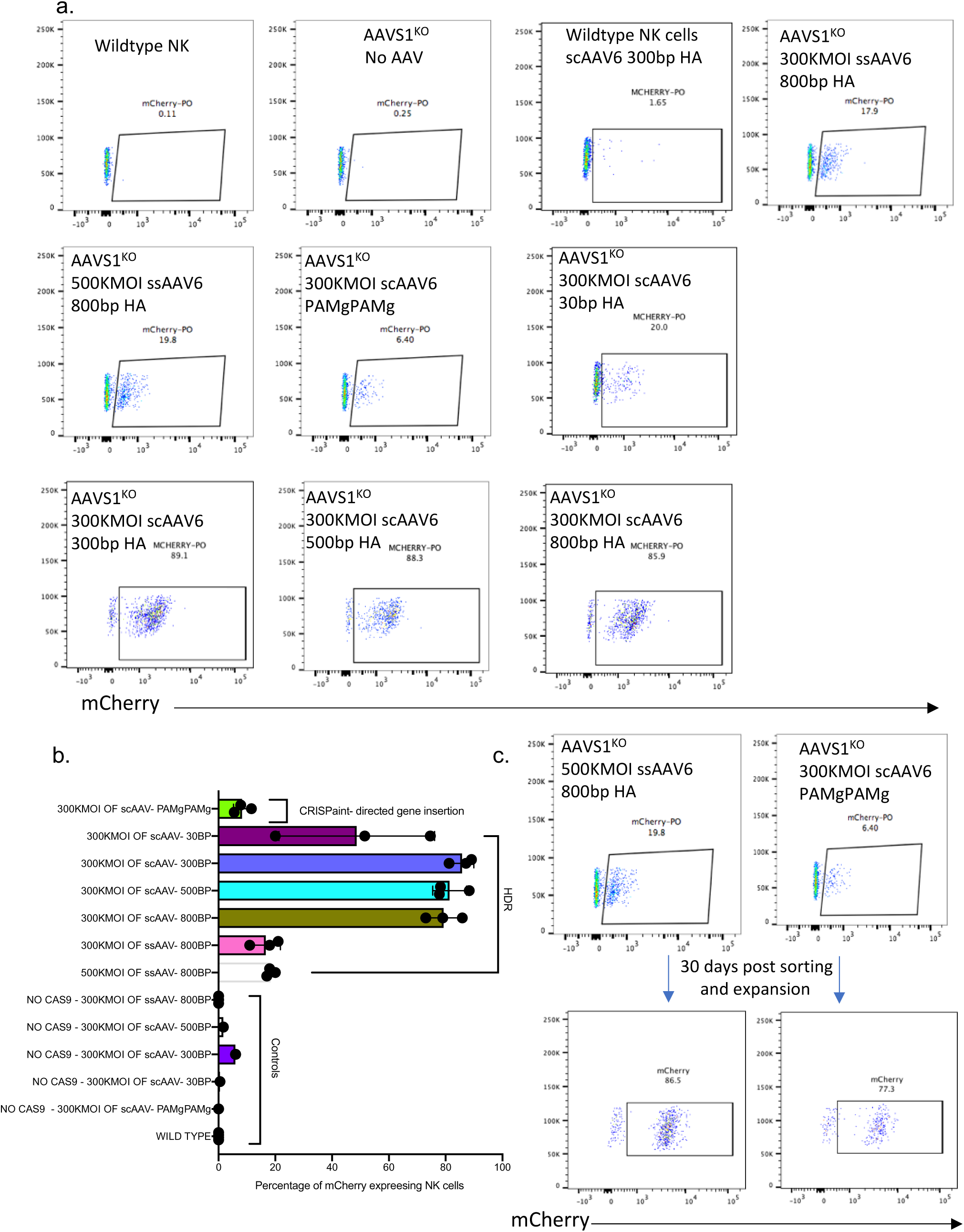
Combinations of AAV6 and Cas9/RNP results in efficient generation of mCherry expressing NK cells. **a,** Representative flow cytometry of human primary NK cells expressing mCherry, 2 days after CRISPR electroporation and AAV6 transduction (MOI = 3 x 10^5^), **b,** Efficiency of Cas9/RNP and AAV6-mediated mCherry expression in human primary NK cells through HR and CRISPaint (n=3). **c,** Stable mCherry expression in NK cells after enrichment and expansion using mbIL21 K562.

### Improved gene insertion by using self-complementary AAV6 and Cas9/RNP

After transduction, scAAV vectors can acquire the necessary double-stranded state in a shorter time frame than ssAAV, which may impact the efficiency of gene insertion. Due to the size limitation of packaging transgenes in scAAV, we designed HAs of varying lengths to minimize the size needed for scAAV backbones. Hence, HAs of 30bp, 300bp, 500bp, and 1000bp length for the right and 30bp, 300bp, 500bp, and 800bp for the left (Figure 2a) were cloned with mCherry into the scAAV backbone and packaged into AAV6 capsid. We then followed the same steps as for the ssAAV, above, to electroporate and transduce the Day 7 expanded NK cells. scAAV with HA ≥ 300bp showed markedly increased efficiency of gene transfer at >80% (Figures 3a and 3b). Stable mCherry gene expression was observed for at least 3 weeks of additional NK cell expansion. When we used the same approach in freshly isolated NK cells, the mCherry expression was significantly lower (1.13% for ss800bp AAV6; 2.9% for sc300bp AAV6, Supplementary Figure 2).

### CRISPaint for gene insertion in NK cells

To overcome the complexity of HAs optimization seen in HDR-directed gene insertion, we tested a homology-independent gene insertion approach called CRISPaint. For the CRISPaint DNA templates, we incorporated double Cas9-targeting sequences of AAVS1 (PAMgPAMg) around the mCherry transgene but within the ITRs of scAAV and packaged it into AAV6 (Figure 2b). Two days after electroporation and transduction we performed flow cytometry to assess mCherry expression in NK cells. The cells which were electroporated and transduced with 300K MOI of scAAV6 delivering CRISPaint PAMgPAMg were found to be up to 6% of mCherry positive. We further sorted and enriched these NK cells and expanded them for 30 days and saw no decline in the percentage that were mCherry positive (Figures 3b and 3c). Although we saw lower efficiency of gene integration using CRISPaint compared to HR-directed gene insertion, this method may still be useful because it allows integration into a user-defined locus without designing homology arms.

### Generation of human primary CD33-CAR NK cells

We designed two CAR constructs comprising the same CD33-targeting scFv, but with a CD4 transmembrane domain and 4-1BB / CD3ζ signaling domain (Gen2) or an NKG2D transmembrane domain and 2B4 / CD3ζ (Gen4v2)^41^ (Figures 4a and 4b). The size of the CAR constructs were too large for suitable packaging into the scAAV backbone, therefore we cloned them into the ssAAV backbone with the largest possible HAs of 600bp. To improve the expression of the CARs, we also incorporated a murine leukemia virus-derived promoter (MND) before the start codon of the CARs instead of the splice acceptor. As with the mCherry vectors, these were packaged into the AAV6 capsid. Seven days after electroporation and transduction, we detected up to 78% CD33 CAR-expressing NK cells (mean 59.3% for Gen2 and 60% for Gen4v2). Of note, the CD33CAR-Gen2 resulted in a higher level of expression on NK cells compared to Gen4v2 (Figures 4c and 4d, Supplementary Figure 3a). We expanded the cells for another week (Day 14) and observed no significant reduction in CAR expression (Figure 4e, Supplementary Figure 3b) or proliferative potential (Figure 4f). Surprisingly, efficient CAR integration (> 60%) was observed with MOIs as low as 10K (Supplementary Figure 4).

**Figure 4.**
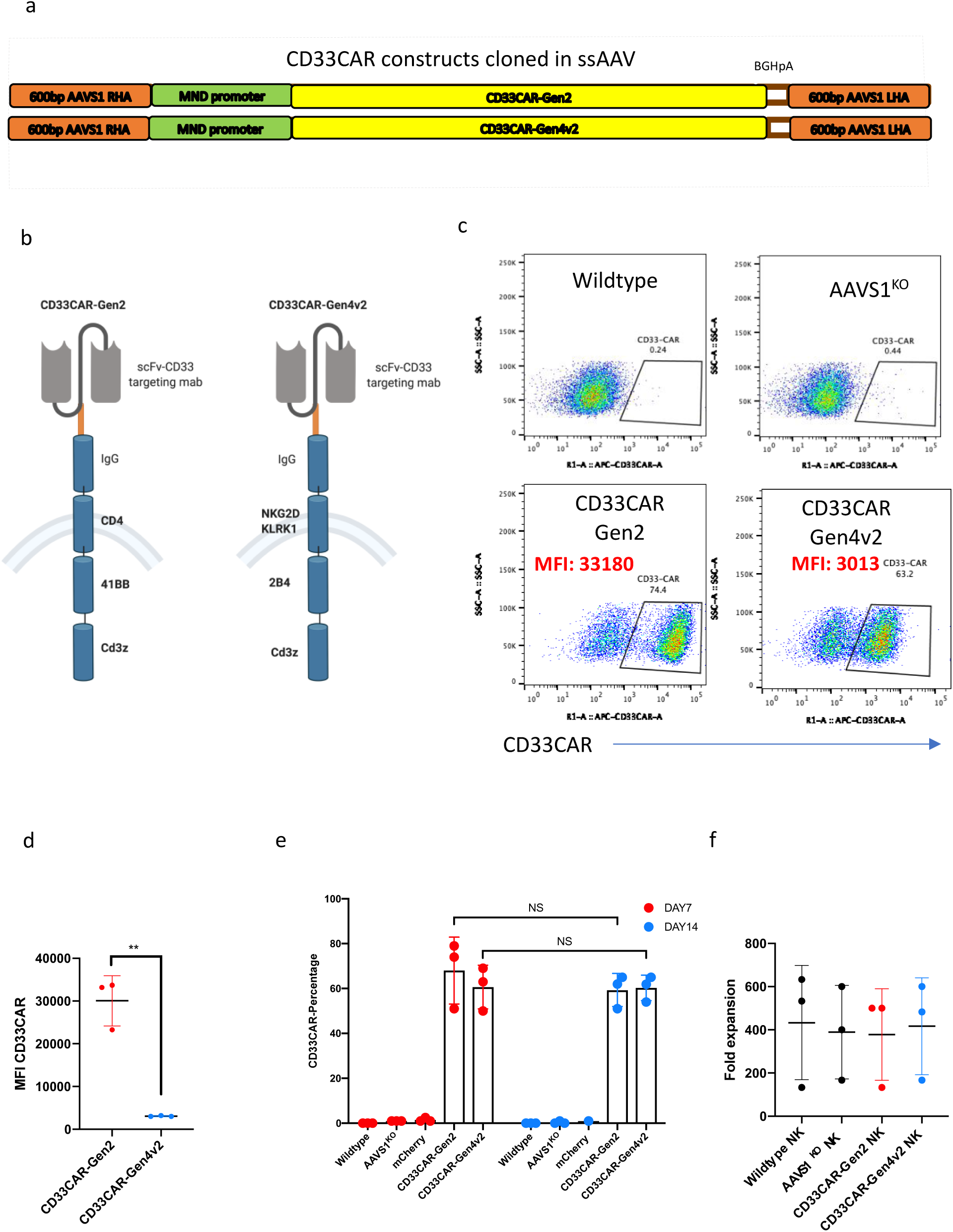
Successful generation of CD33CAR expressing NK cells using combination of Cas9/RNP and AAV6. **a and b**, Schematic of anti-CD33 CAR constructs (Gen2 and Gen4v2) designs with HAs for AAVS1 targeting site and cloned in ssAAV. **c** Representative flow cytometry showing the expression level of CD33CAR on NK cells, 7 days after Cas9/RNP electroporation and AAV6 transduction (MOI = 3 x 105). **d,** MFI of CD33CAR expression of Gen2 was significantly higher than Gen4v2, **P = 0.0014, measured by unpaired t-test. **e,** CD33CAR expression level on NK cells seven and fourteen days after transduction and electroporation showed no significant reduction (n=3). **f,** Fold expansion of CD33CAR expressing NK cells on feeder cells for 14 days starting from 3 x 10^5^ cells (n=3) was similar to wildtype NK cells. n.s are from multiple comparison 2way ANOVA.

### Detection of the transgene at the targeted AAVS1 locus and unintended insertion sites

Using PCR with primers to flanking and inter-transgenic regions (Figure 5a, Supplementary Table 2), we confirmed the DNA integration of the transgenes (Figure 5b). Additionally, targeted locus amplification (TLA) was used for whole-genome mapping of CD33CAR-Gen2 integration in CAR-expressing NK cells with a sensitivity of detecting random integrations of more than 5%, which demonstrated high prevalence of vector integration at the targeted location in chromosome 19, with low level random integrations identified throughout the genome (Figure 5c, Supplementary Figure 5, Supplementary Table 3). There was no indication of a dominant secondary off-target integration site. One sequence variant (Supplementary Table 4) and four structural variants (Supplementary Figure 6, Supplementary Data 1) were detected.

**Figure 5.**
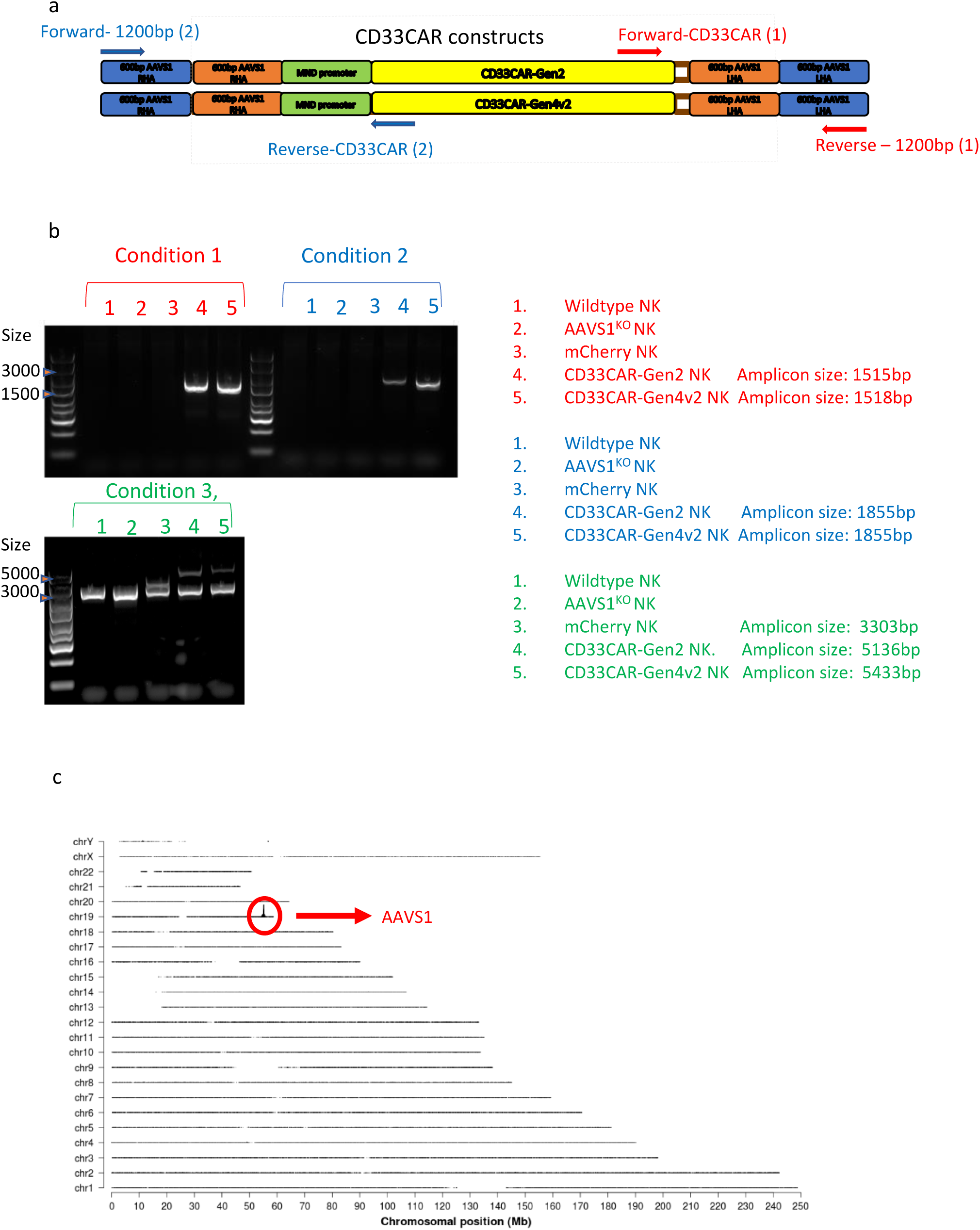
Integration of the transgene in AAVS1 locus was confirmed by PCR and TLA. **(a),** Schematic of PCR primers designed inside and outside of CD33CARs encoding DNA and integrated in AAVS1. **(b),** Amplicons were amplified and visualized on 1% agar gel only in NK cells with successful CD33CAR gene insertion at AAVS1 locus (condition 1 and 2). The gene insertion in human primary NK cells also was seen when primers designed outside of the transgenes and were used to amplify AAVS1 locus in wildtype, mCherry or CD33CARs (condition 3, primers: Forward-1200bp (2) Reverse – 1200bp (1)), Supplementary Table 2. **(c)**, TLA sequence coverage across the human genome using designed primers to detect integration of CD33CAR-Gen2 in day 14 cells. The chromosomes are indicated on the y-axis, the chromosomal position on the x-axis. Identified integration site is encircled in red. The, Supplementary Table 3 indicates the primers used for the TLA sequencing

### Human primary CAR-NK cells have enhanced antitumor activity

To determine whether the CD33CAR enhanced NK cell killing of AML cells, we performed Calcein AM based cytotoxicity assay with two CD33-expressing AML cell lines (Kasumi-1 and HL60) and one patient-derived sample (AML10) (Supplementary Figure 7) using CD33CAR-NK cells generated from three different healthy individuals. CD33CAR-gen2 and CD33CAR-gen4v2 NK cells showed a significantly higher degranulation and target cell lysis when cocultured with Kasumi-1 or HL60 cell lines in comparison to wildtype or *AAVS1^KO^* NK cells. (Figures 6a-6f). Importantly, we showed significantly higher antitumor activity of CD33CAR NK cells against *AML-10,* a primary human AML sample derived from a patient with relapsed and refractory AML (Figures 6g and 6h)^5, 19^. Overall, CD33CAR-Gen2 NK showed better cytotoxicity in comparison to CD33CAR-Gen4v2 NK cells. Using real-time assessment of cytotoxicity (xCELLigence), we showed that CD33CAR-NK cells kill the CD33-expressing AML cells more completely and with faster kinetics than the same donor WT NK cells (Figure 6f).

**Figure 6.**
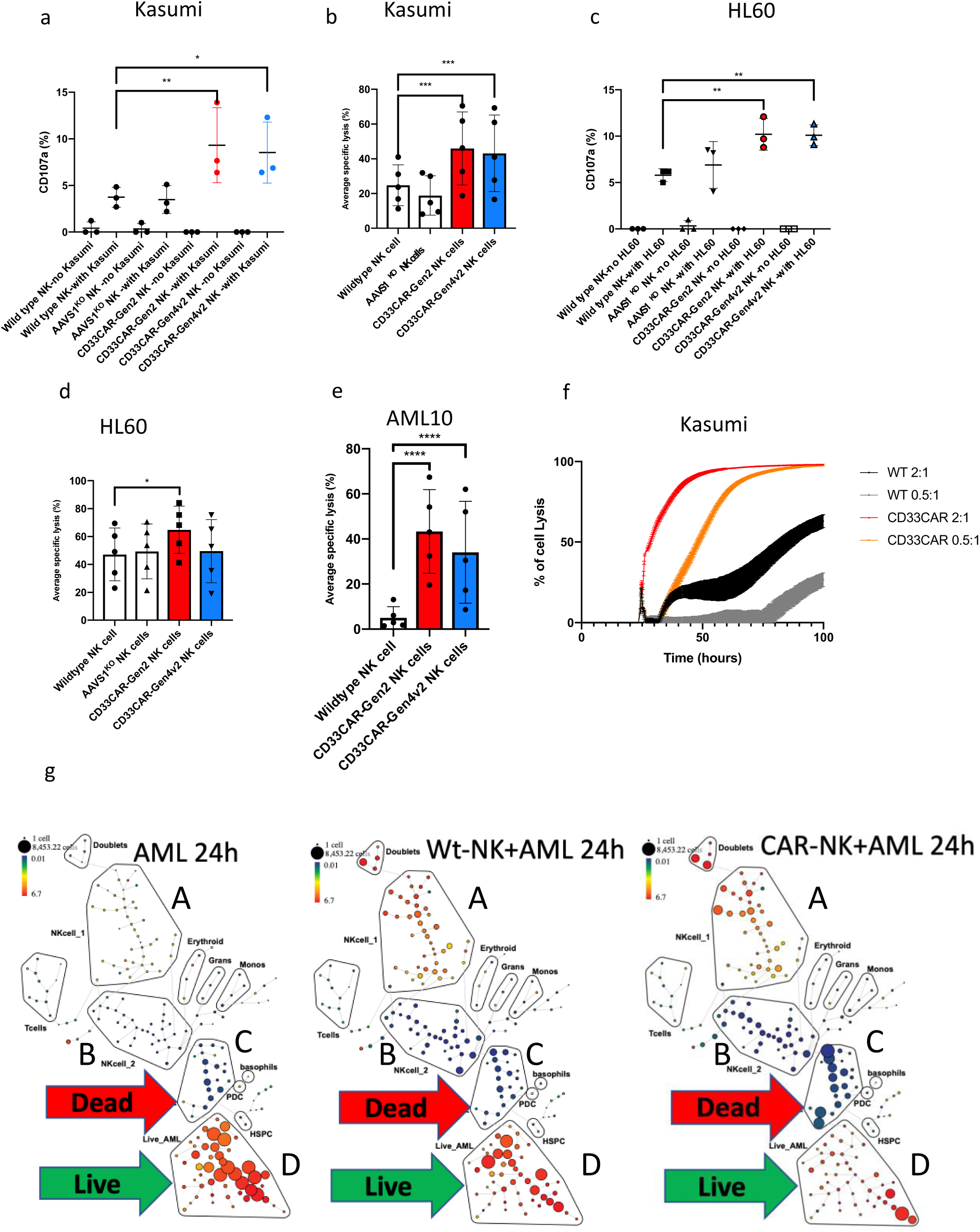
CD33CAR NK cells have enhanced anti-AML activity. CD33CAR NK cells degranulate significantly higher than wildtype NK cells when cocultured with *Kasumi-1, \*\** adjusted P value= 0.004 **(a)** and show higher killing ability **(b).** The enhanced degranulation **(c)** and killing against *HL60*, * adjusted P value= 0.01 **(d)** as shown in representative cytotoxicity assay performed in different effector:target ratios **(b and d)** and in three donors, **** adjusted P value <0.0001., * adjusted P value= 0.01. CD33CAR-Gen2 and Gen4v2 significantly killed higher *AML-10* primary cells, **** adjusted P value <0.0001 **(e).** P Values are from multiple comparison 2way ANOVA. CD33CAR-Gen2 NK cells also showed faster killing of AML cells in compare to WT-NK cells in a xCelligence killing assay **(f).** We also observed the specificity and killing potency of CD33CAR-NK cells against CD33-expressing primary AML cells in CyTOF assay **(g).**

### Mass Cytometry showed enhanced killing and specificity of CD33CAR-NK cells against AML

After co-culture of WT or CD33-CAR NK cells with primary AML, we could identify four main populations (Figure 6g) by both manual gating and SPADE clustering: live proliferating NK cells, quiescent NK cells, live proliferating AML cells, and dying AML cells. At 3 hours, control primary AML cells were ∼66% viable, whereas viability decreased to 56% when co-cultured with WT-NK cells and only 21% when co-cultured with CD33CAR NK cells. At 24 hours, control primary AML cells recovered to 89% viability, compared to 76% when co-culture with WT-NK cells and only 42% when co-cultured with CD33CAR NK cells. At 3 hours, the surviving AML cells had a 20-fold reduction in CD33 expression when cultured with the CD33CAR NK cells (median of 104 counts down to 5 counts) while there was minimal change in CD33 expression in the WT NK cell co-culture (median of 104 counts down to 87 counts). This difference persisted at 24 hours, at which time the median CD33 counts were 25 for CD33CAR NK cells, 118 for WT NK cells, and 120 in control AML without NK cells. Co-culture with AML also increased NK activation markers (CD69, CD99, CD71, NKG2D, CD16, and CD45) at 24 hours compared to the NK cells cultured alone. The CD33CAR NK cells had lower levels of activation markers at baseline that increased more with co-culture, Together, these data show that CD33CAR-NK cells specifically target CD33-expressing AML and are more activated by the AML targets compared to WT-NK cells (Figure 6g).

## Discussion

Gene modification in primary human NK cells has always been challenging; here, we report a successful, highly efficient site-directed gene integration into human primary NK cells using a combination of electroporation of Cas9/RNP and single-stranded or self-complementary AAV6 gene delivery through HR and homology-independent gene insertion (CRISPaint). Here for the first time, we showed how the expression level of genes regulating HR and NHEJ pathways in human NK cells alter during expansion with FC21 and provides an optimal condition for site-directed gene insertion. We also demonstrated that AAVS1 could host and express exogenous genes in a very highly efficient level, as shown previously in T cells and NK cells^10^. Furthermore, we showed that a range of HAs from 30-1000bp that can be used for gene insertion into the AAVS1 locus in NK cells, but that the shortest optimal length is at 300bp when used in scAAV6. This helps researchers to choose an optimal HA based on the size of their exogenous DNA that they will introduce in NK cells. CRISPaint gene insertion can be potentially used for tagging endogenous genes and be used for studying the biology of proteins in NK cells.

Transcripts that are delivered via AAV vectors can be packaged as a linear single-stranded (ss) DNA with a length of approximately 4.7 kb (ssAAV) or as linear self-complementary (sc) DNA (scAAV). The benefit of the scAAV vector is that it contains a mutated inverted terminal repeat (ITR), which is required for replication and helps to bypass rate-limiting steps of second strand generation in comparison to ssDNA vectors^42^. Due to the limitation in the packaging capacity of scAAV, we designed 30bp, 300bp, 500bp, and 800-1000 bps of HAs for the right and left side of the Cas9-targeting site to find the most optimal length of HAs and to provide possible lengths of HAs to be chosen based on the size of transgenes by researchers (Figure 2a). Additionally, due to limitations in packaging capacity compared to ssAAV, scAAV is not suitable for larger transgenes such as chimeric antigen receptor (CAR) targeting CD33^42^. Therefore, based on the size of transgenes, we designed and tested both ssAAV and scAAV, which provides a wide range of options for gene insertion in primary NK cells.

Since designing homology arms is a time-consuming procedure and requires multiple optimizations, we also investigated the CRISPaint approach, a homology-independent method for gene insertion or tagging. In this method, the same Cas9 targeting site, including the crRNA and PAM sequence, is provided in the DNA template encoding the gene of interest. Upon the introduction of the Cas9 complex, both template and genomic DNA will be cut simultaneously. As a result, the CRISPaint template will be presented as a linearized double-stranded DNA that can be integrated through non-homology repair machinery (Figure 2b)^14, 31^. We also used the combination of Cas9/RNP and AAV6 gene delivery and generated two different human primary CD33CAR NK cells with enhanced anti-AML activity. Our results also showed that the gene-modified NK cells could be subsequently expanded with FC21, enabling the production of large numbers of gene-modified NK cells for cancer immunotherapy. Overall, our novel method can be used for several applications in immunology, cancer immunotherapy, and studying the biology of NK cells.

## Material and Methods

### Human NK Cell Purification and Expansion

NK cells were purified as previously described^11^. Briefly, NK cells were isolated from PBMC collected from healthy individuals using RosetteSep™ Human NK Cell Enrichment Cocktail. Purified NK cells were phenotyped using flow cytometry as >90% CD3-negative/CD56-positive population (Supplementary Figure 1a). These cells were then stimulated with irradiated feeder cells (FC21) comprised of K562 transduced with 4-1BBL and membrane-bound IL-21 (CSTX002)^17^ at a ratio of 2:1 (feeder: NK) as previously described ^12, 18^ ^19^. The stimulated cells were cultured for 7 days in serum-free AIM-V/ICSR expansion medium containing 50 IU/mL of IL-2^20^.

### ATAC-seq assay

Freshly-isolated (naïve) and FC21-expanded NK cells from two donors were cryopreserved in aliquots of 100,000 viable cells/vial before processing for ATAC-seq. ATAC-seq was performed as previously described^21^. DNA libraries were sequenced using Illumina HiSeq 2500 at 50 bp paired-end reads.

### Cas9/RNP electroporation for targeting AAVS1 in NK cells

AAVS1 was targeted using gRNA (crRNA: 5’GGGGCCACTAGGGACAGGAT) via electroporation of Cas9/RNP into Day 7 expanded NK cells as described before^11^. Briefly, 3 x 10^6^ expanded NK cells were harvested and washed twice with 13ml of PBS followed by centrifugation for 5 minutes at 400g and aspiration of PBS. The cell pellet was resuspended in 20ul of P3 Primary Cell 4D-Nucleofector Solution. 5ul of pre-complexed Cas9/RNP (Alt-R® CRISPR-Cas9 crRNA, Alt-R® CRISPR-Cas9 tracrRNA, and Alt-R® S.p. HiFi Cas9 Nuclease V3) (Integrated DNA Technologies, Inc., Coralville, Iowa), targeting AAVS1 and 1ul of 100uM electroporation enhancer (Alt-R® Cas9 Electroporation Enhancer) were added to the cell suspension. The total volume of 26ul of CRISPR reaction was transferred into 4D-Nucleofector^TM^ 16-well Strip and electroporated using program EN-138 (Supplementary Figure 1a). After electroporation, the cells were transferred into 2ml of media containing 50IU of IL-2 in a 12 well plate and incubated at 37 degrees and 5% CO2 pressure. Two days post electroporation, cells were stimulated with 2 x 10^6^ feeder cells, and 8ml fresh media complemented with 50IU was added in cell suspension and kept in a T25 flask.

### Inference of CRISPR Edits (ICE) mutation detection assay

To measure the indel rate in AAVS1^KO^ NK cells, the Cas9/RNP targeted site was PCR amplified using forward and reverse primers described in Supplementary Table 1. The amplicons were sequenced using Sanger sequencing, and results were analyzed using ICE ^22^ (Synthego, Menlo Park, CA).

### RNA-seq sample preparation and sequencing

Total RNA was purified from naïve resting, expanded resting, naïve IL21-stimulated, and Day 14 FC21-expanded NK cells using the Total RNA Purification Plus Kit (Norgen Biotek, Ontario, Canada). The resulting total RNA was sequenced and analyzed as described before ^12^.

### AAV6 production

The transgenes cloned into ssAAV or scAAV plasmids were packaged in AAV6 capsids as described before^23^.

### Combining Cas9/RNP and AAV6 to generate mCherry and CAR NK cells

A media change and resuspension at 5 x 10^5^ cells per ml was performed on day 6 of NK cell expansion one day before experimental manipulation. The NK cells were electroporated with Cas9/RNP targeting AAVS1 on day 7, as described above. Thirty minutes after electroporation, 3 x 10^5^ live cells were collected and resuspended at 1 x 10^6^ cells per ml in media containing 50IU IL2 (Novartis) in a 24 well plate in a total volume of 300ul. For each transduction condition with ssAAV6 or scAAV6 to deliver HR or CRISPaint DNA encoding mCherry or CD33CARs, we transduced 3 x 10^5^ electroporated cells with 300K MOI (10-500K MOI if needed). Negative controls included as NK cells that were not electroporated, or were electroporated with Cas9/RNP but not AAV transduced, or were transduced with 300K MOI of AAV6 without electroporation of Cas9/RNP. The day after electroporation and transduction, we added 300ul of fresh media containing 50IU of IL2 to each well without changing the old media. The cells were kept in culture for 48 hours after electroporation and were then restimulated with 2 x 10^6^ feeder cells and kept in a total volume of 2ml media containing 50IU in 12 well plate, without changing the old media. 48 hours later, 8ml fresh media supplemented with IL2 was added to cells, a total volume of 10ml was kept in a T25 flask. At day 7 post-transduction, cells were re-stimulated with feeder cells at a ratio of 1:1 and grown for one more week, every 2 days fresh media was added to the cells.

### Flow Cytometry for detection of CAR-NK cells

7 days and 14 days following electroporation, 5 x 10^5^ NK cells were washed twice with staining buffer containing 2% FBS in PBS. Next, 2.5ug of recombinant human siglec-3/CD33 Fc chimera protein, (CF; R&D systems #1137-SL-050) was added to cell suspension in a total volume of 80ul and incubated for 30 minutes at 4C. Cells were washed twice with staining buffer before staining with 2ul of Alexa Fluor® 647 affinipure goat anti-human IgG, Fcγ fragment specific, (Jackson ImmunoResearch #109-605-098) at 1:100 ratio in 200ul of staining buffer and kept at 4C for 30 minutes. Once stained, cells were washed twice with staining buffer then acquired on MacsQuant flow cytometers. Flow cytometry data were analyzed using FlowJo software (FlowJo, LLC).

### Cytotoxicity assay

Cytotoxicity assays were performed for 3-4h as described previously using a calcein-acetoxymethyl-release assay^19^. Cytotoxicity was assessed against Kasumi-1, HL60, or AML10 cells at different effector:target ratios as defined in Figure 6a-6e.

### CD107a staining

NK cells and cancer cells were cocultured at 10:1 ratio and supplemented with 20ul of PE mouse anti-human CD107a antibody (BD Pharmingen™, #555801) in a total volume of 220ul in a 96 well plate at 37C incubator for 90 minutes. The cells were washed with staining buffer once and analyzed on MacsQuant flow cytometer.

### PCR-based detection of transgenes integration

In-out PCR was performed using 2 pairs of primers (Figures 5a and 5b and Supplementary Table 2) designed inside or outside of the CD33CAR constructs. We also added a set of primers to amplify 1200bp right and left flanking region of Cas9 targeting and transgene integration site (Figure 5a). PCRs were performed using the Platinum™ Taq DNA polymerase high fidelity kit (Thermofisher #11304011).

### Targeted Locus Amplification (TLA)

For the whole-genome mapping of CD33CAR-Gen2 integration, we used the TLA technology (Cergentis B.V.)^24^. The genomic DNA from CD33CAR-expressing NK cells was isolated using Qiagene DNeasy Blood & Tissue Kit and crosslinked, fragmented, and re-ligated using the kit provided by Cergentis, then submitted to Cergentis for sequencing^24^.

### Real-time potency assessment for suspension target cells killed by WT and CD33CAR NK cells

The xCELLigence RTCA MP instrument (ACEA Biosciences) was utilized as previously described^25^. Briefly, the CD29 tethering reagent was coated on the plate (2 μg/mL, for 3 hours at 37°C) to immobilize the Kassumi cell line, seeded at 6 x 10^4^/well. After 20-24 hours, WT or CD33CAR NK cells were added at indicated E:T ratios. An “effector cell only” control was included in the presence of tethering reagent. Cell Index values reflecting viable target cell adherence were applied to xIMT software to plot percentage cytolysis.

### AML-NK Cell Co-Culture for Mass Cytometry

Cells were grown in SFEM II (StemCell, Cambridge, MA) supplemented with the following cytokines; SCF, IL-6, TPO, FLT3, GM-CSF, G-CSF, IL-3 (20ng/ml) and EPO (10ng/ml). A de-identified primary AML sample was obtained from OSU Leukemia Tissue Bank consistent with Declaration of Helsinki. NK or CAR-NK cells at a target:effector ratio of 1:2 were cultured for either 6 or 24 hours. Following culture, cells were prepared for staining as previously described^26^.

### Mass Cytometry Staining and Analysis

Fixed cells were washed with CSM as previously desrcibed^27^. Antibodies used for this study are listed in Supplemental Table 5. Samples were resuspended in 1:20 dilution of four elemental equilibration beads (Fluidigm) at a concentration of 1 million cells/ml. FCS generated files were normalized using a normalization tool developed by Finck et al and analyzed on Cytobank (www. cytobank.org)^28, 29^.

Once the singlet gate was established, cells were identified using markers and analyzed using SPADE^30^.

## Acknowledgments

This work was supported by funding from the CancerFree KIDS Pediatric Cancer Research Alliance (M.NK), Hyundai Hope on Wheels Foundation (D.A.L), and NIH-NCI (U54-CA232561, D.A.L). We thank the Institute for Genomic Medicine at Nationwide Children’s Hospital for their assistance with nucleic acid sequencing. We also aknowledge Yasmein Sezgin, for her assistant in performing some of the experiements.

---

**Supplementary Figure 1.**
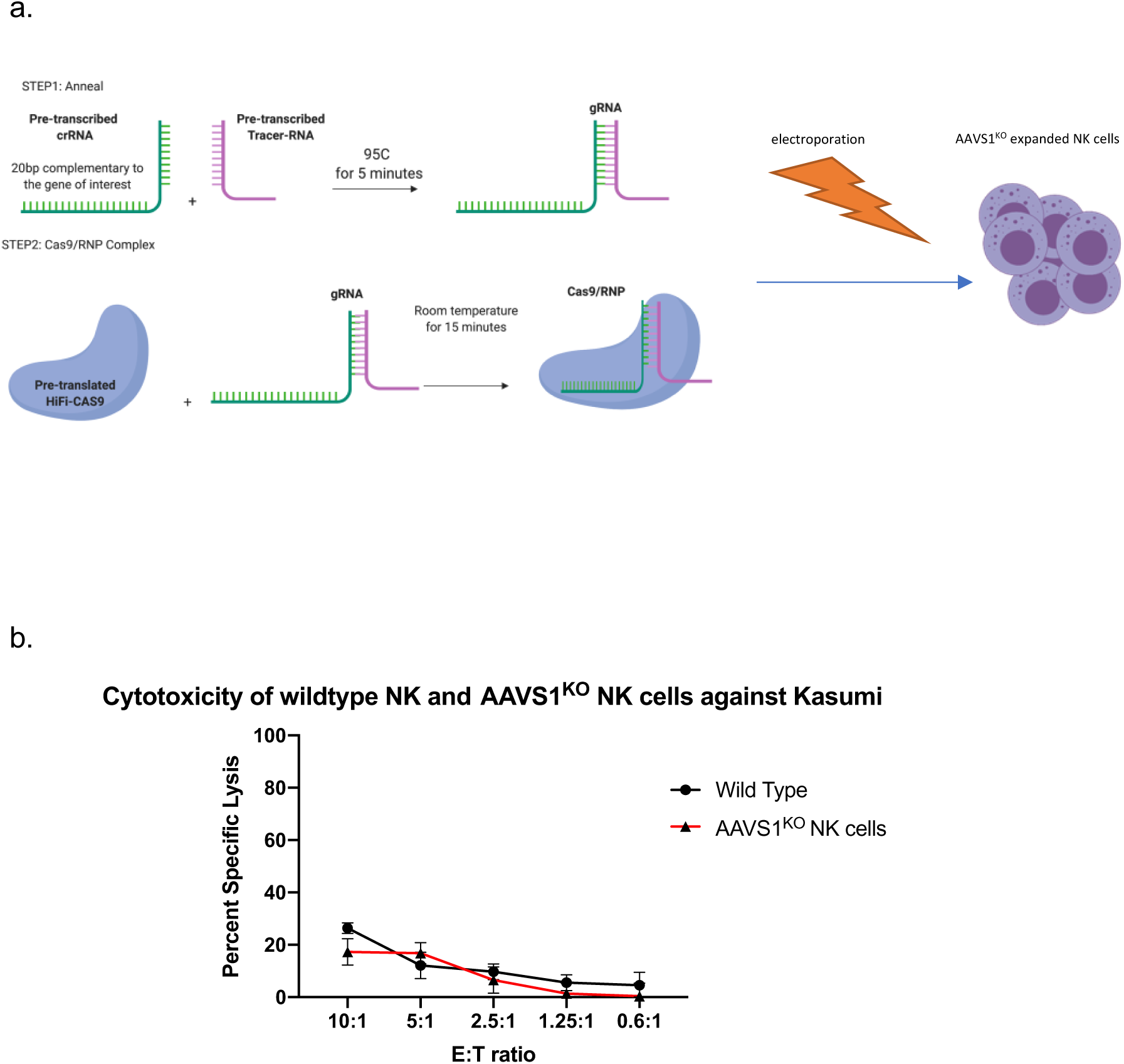
Targeting AAVS1 in expanded CD3^negative^ CD56^positive^ NK cells does not alter normal function of the cells. **a,** Schematic of workflow for electroporation of Cas9/RNP into day 7 expanded human primary NK cells to target AAVS1. **b,** Cytotoxicity assay of AAVS1^KO^ NK cells does not show any suppression in their antitumor activity againts AML cell line.

**Supplementary Figure 3.**
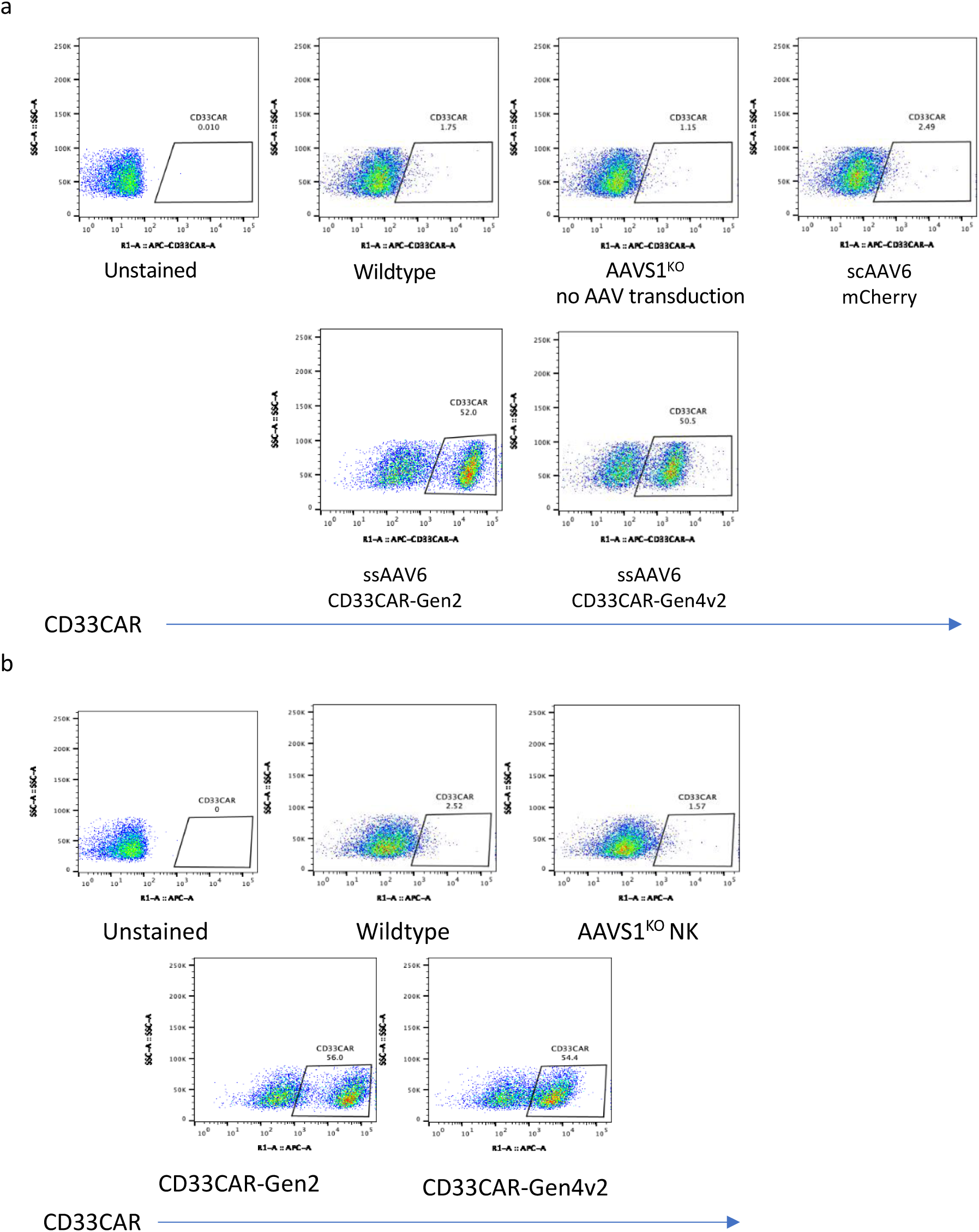
Representative flow cytometry analysis of CD33CAR expression level 7 days **(a)** and 14 days **(b)** post electroporation and AAV6 transduction in human NK cells.

**Supplementary Figure 4.**
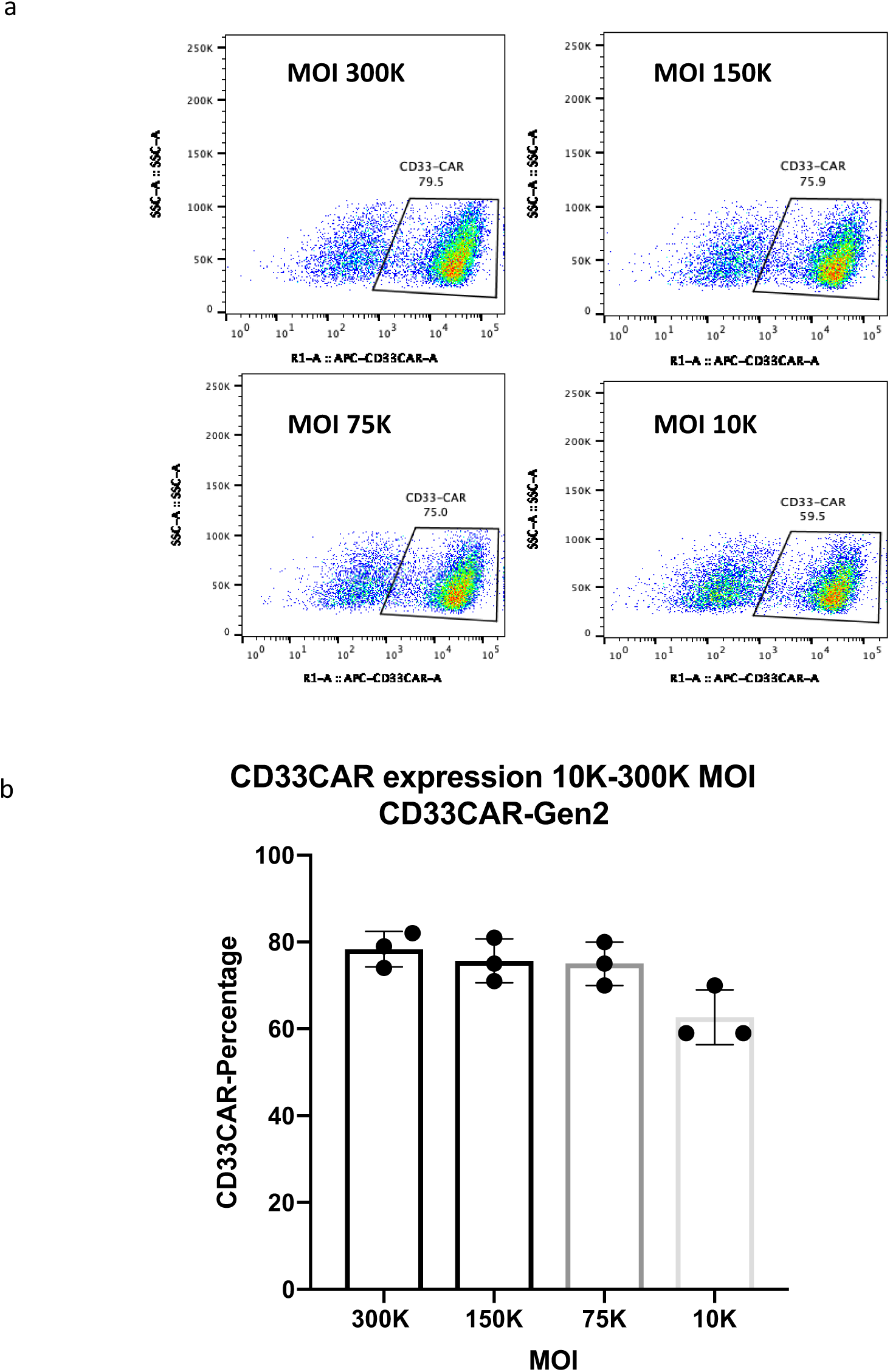
Representative flow cytometry **(a)** analysis of CD33CAR-Gen2 expression level in NK cells transduced with 10K-300K MOI of ssAAV6 encoding CD33CAR-Gen2 showed successful expression of CAR on NK cells isolated from three ealthy donors **(b)**.

**Supplementary Figure 5.**
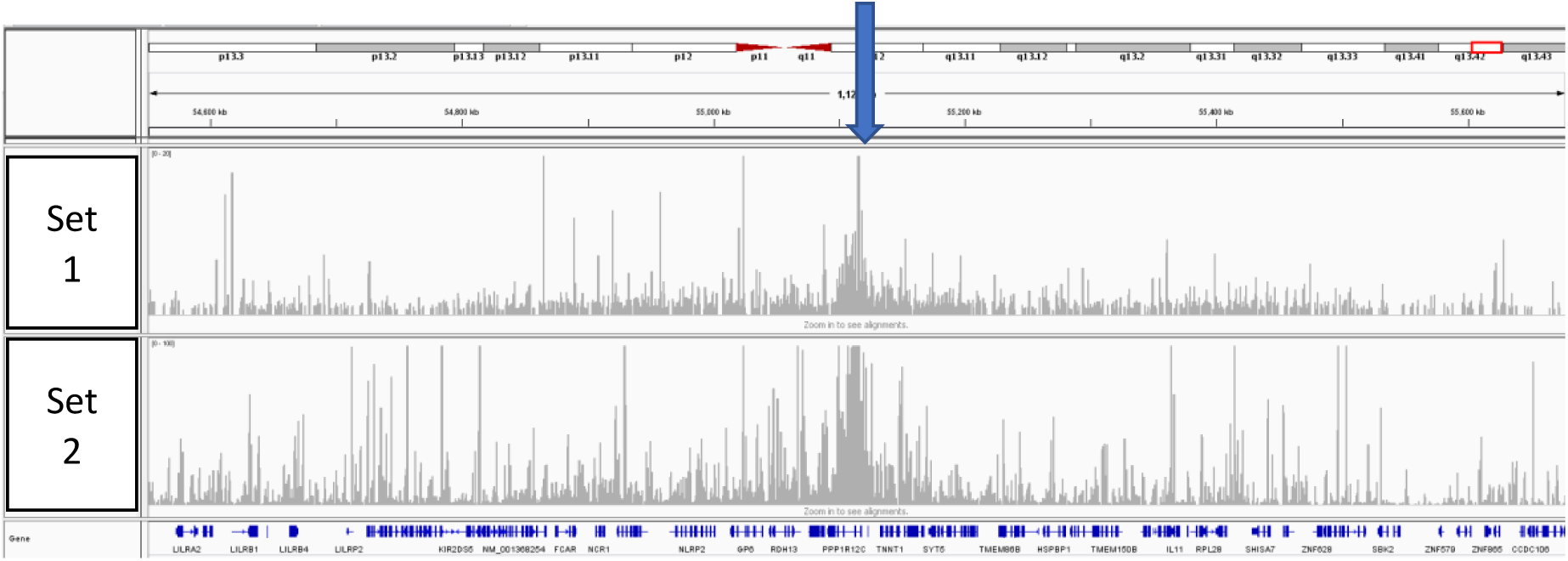
TLA sequence coverage (in grey) across the vector integration locus, human chr19:54,550,476-55,682,266. The blue arrow indicates the location of the breakpoint sequences. Y-axes are limited to 20x and 100x resp. The coverage profile this figure shows that no genomic rearrangements have occurred in the region of the integration site. From this data it is concluded that the vector has integrated as intended in human chromosome chr19: 55,115,754-55,115,767. According to the RefSeq this is in intron 1 of PPP1R12C. Other integration sites were observed between chr19: 55,115,155-55,116,371. According to the RefSeq this is also in intron 1 of PPP1R12C.

**Supplementary Figure 6.**
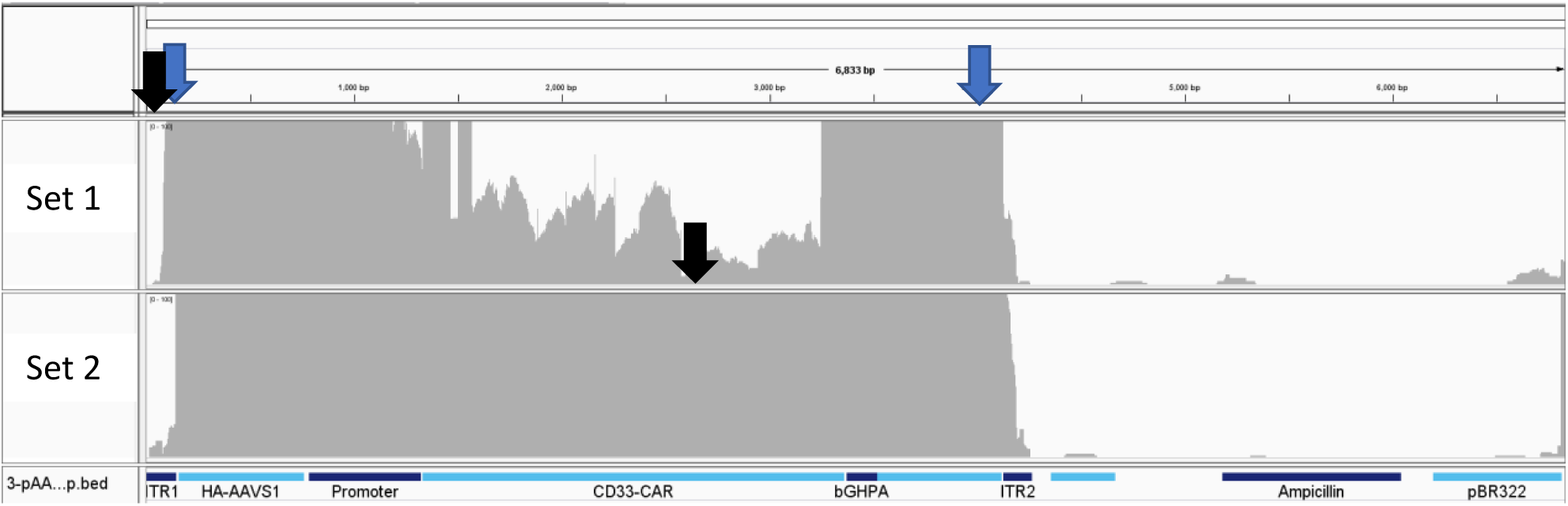
NGS sequencing coverage (in grey) across the vector. Black arrows indicate the primer location. The blue arrows indicate the locations of the identified vector-genome breakpoint sequences (described below). The vector map is shown on the bottom. Y-axes are limited to 100x.

High coverage is observed across the region between the ITR sites, vector sequence Vector: 12-4,255. Low/no coverage is observed across the Vector: 0-11 and 4,256-6, 864 indicating the backbone has not integrated in a large proportion of this sample, potentially a small subset of the sample might contain the backbone as well. Also, coverage is observed at the ITRs, indicating that next to the integration through the homology arms also ITR based integrations occurred in the sample. Sequence variants and structural variants were called in the covered regions.

## Sequence variants

A single sequence variants of the transgene was detected by this method as shown in Supplementary Table 4. The frequency of detection suggests this variant was present within the AAV6 vector itself.

**Supplementary Figure 7.**
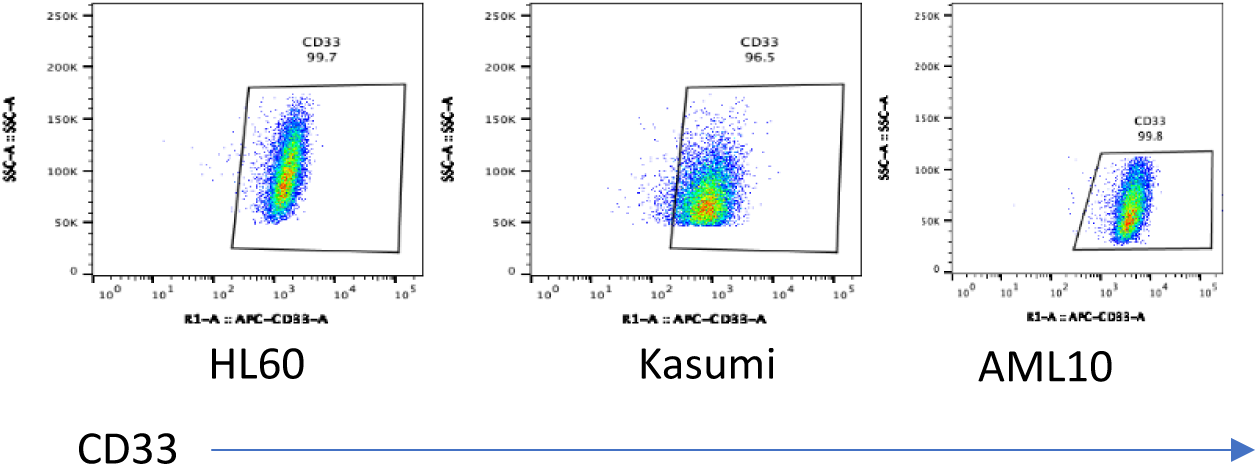
CD33 expression level in different cancer cells.

**Supplementary Figure 8.**
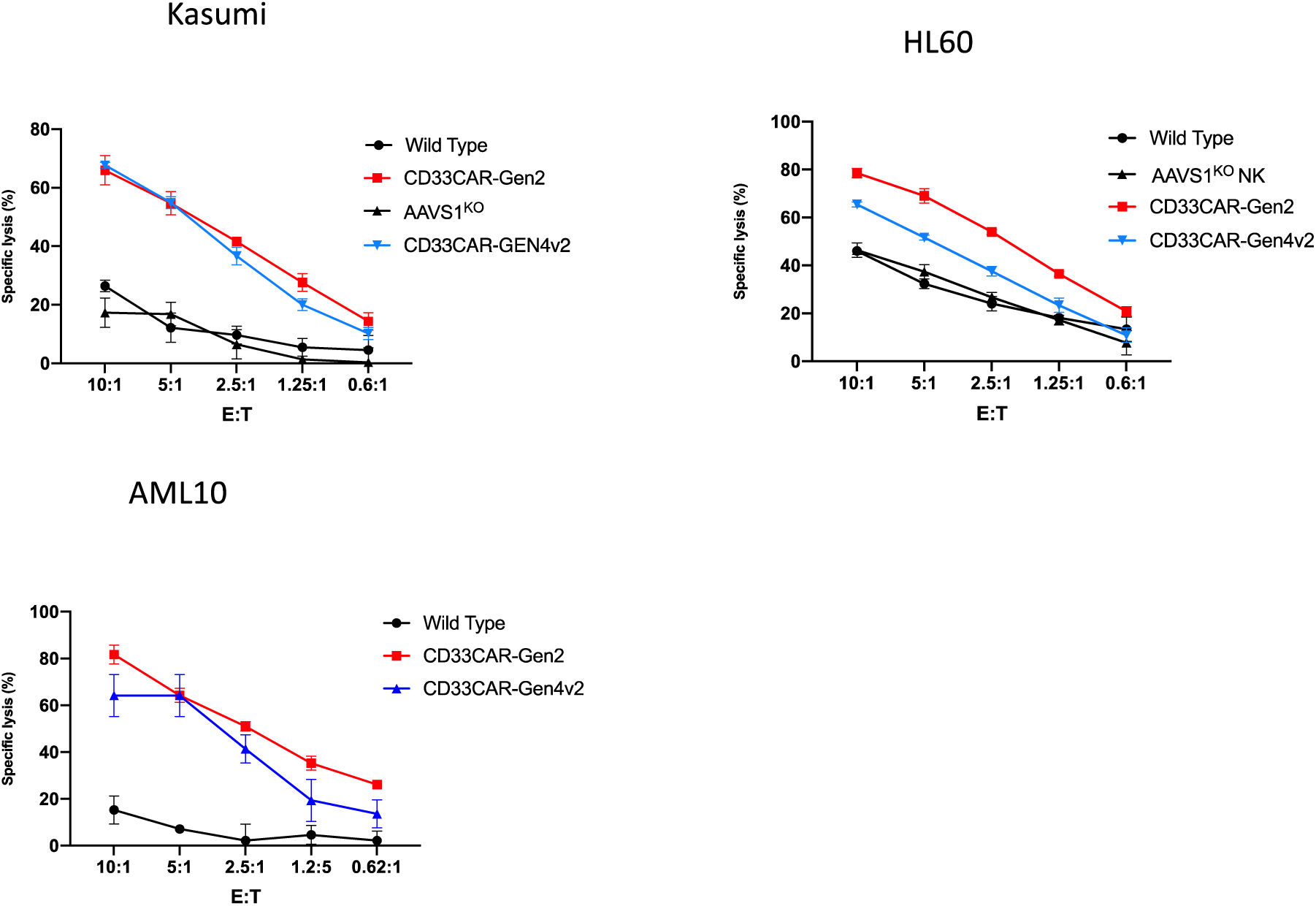
Expressing CD33CAR on NK cells also enhances antitumor activity of NK cells against *Kasumi-1* as shown in representative cytotoxicity assay performed in different effectoπtarget ratios This enhanced cytotoxic activity was observed against *HL-60* only in CD33CAR-Gen2 NK cells.

**Supplementary Table 1:**
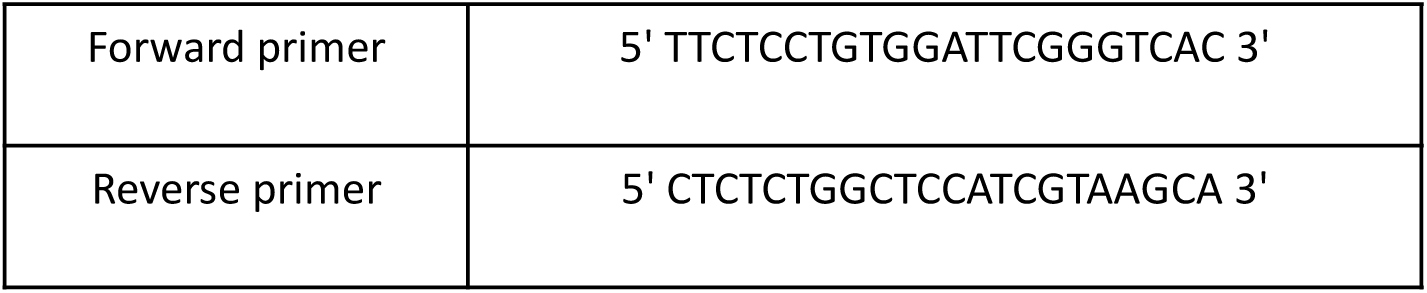
Primers used for Inference of CRISPR Edits (ICE) mutation detection assay

**Supplementary Table 2:**
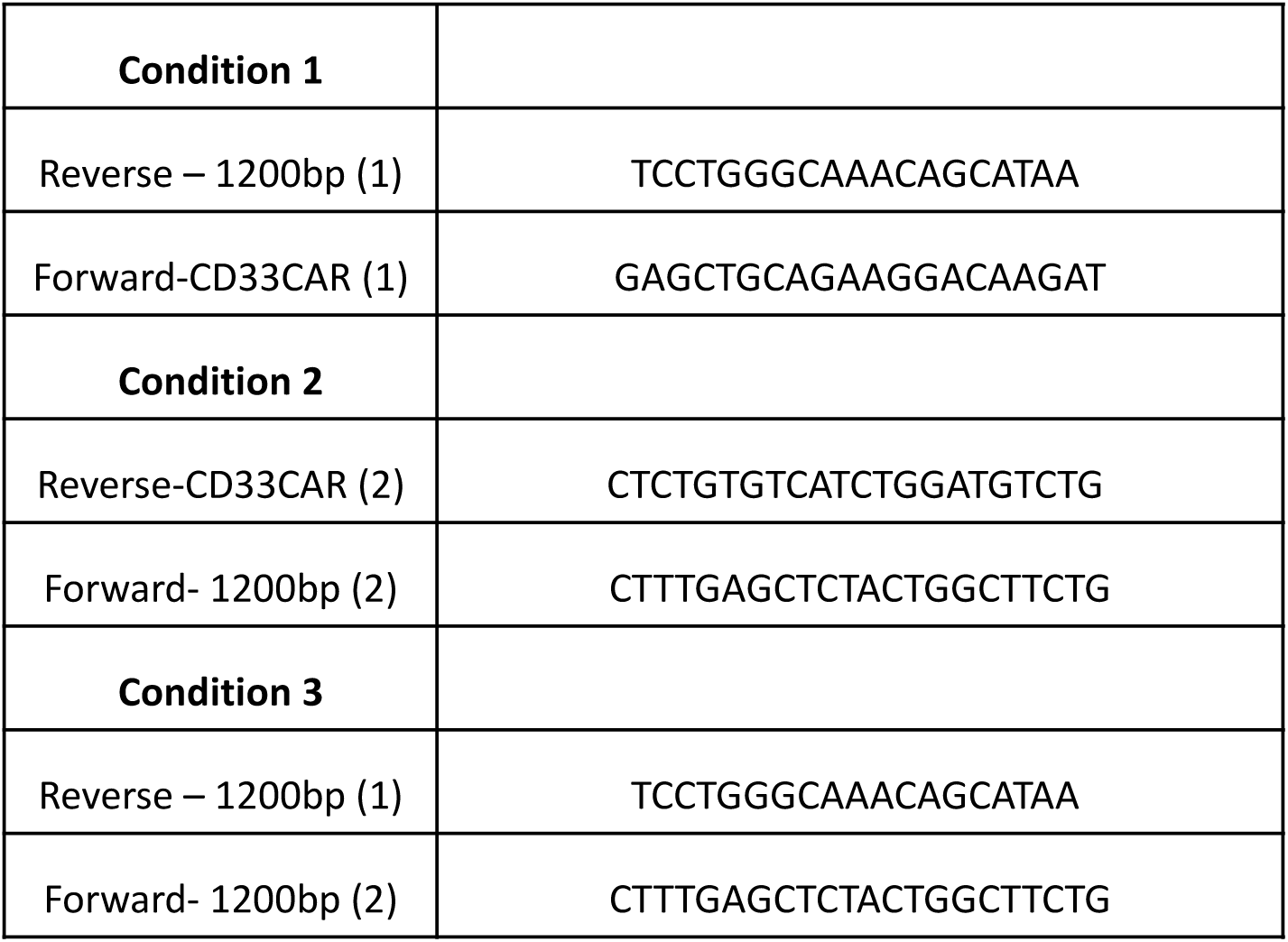
Primers used for PCR-based detection of transgenes integration

**Supplementary Table 3:**
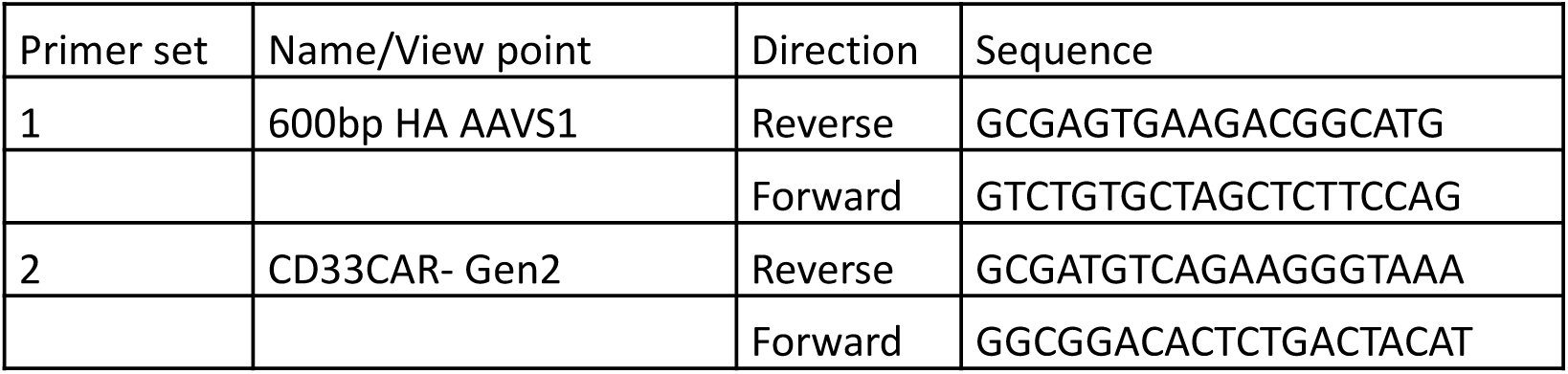
Vector-specific primer sets used for targeted locus amplification (TLA).

**Supplementary Table 4:** Transgene sequence variant identified

|  |  |  |  | Primer set 1 |  | Primer set 2 |  |
| --- | --- | --- | --- | --- | --- | --- | --- |
| Region | Position | Reference | Mutation | Coverage | % | Coverage | % |
| 600bp HA<br>AAVS1 | 4,116 | T | C | 193 | 37 | 341 | 14 |

## Supplementary Data 1

### Identifying structural variants

4 vector-vector breakpoints were found. All fusions were located at the annotated homology arm. Due to the heterogeneous nature of the sample it is expected that these fusions are only present in a subset of the sample. It should be noted that three out of four fusions show 9-12 bp homology which might indicate technical bias.

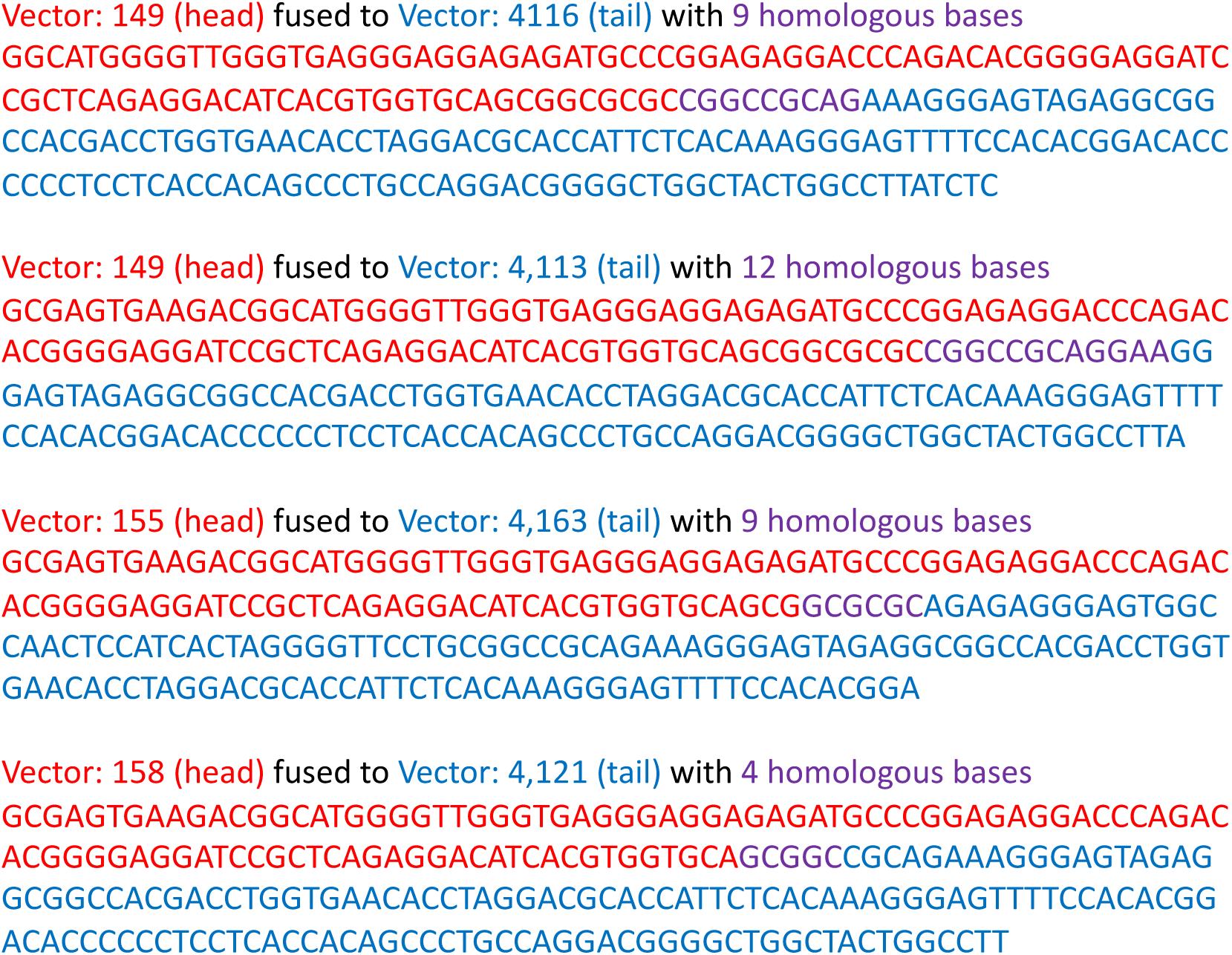

**Supplementary Table 5:** List of the antibodies used for CyTOF analysis

|  |
| --- |
| 140Ce_cPARP |
| 151Eu_CD123 |
| 153Eu_HLA-DR |
| 154Sm_CD69 |
| 156Gd_CyclinB1 |
| 165Ho_pRb |
| 175Lu_Perforin |
| 142Ce_CD7 |
| 148Nd_CD34 |
| 152Sm_CD33 |
| 167Er_CD99 |
| 139La_CD41 |
| 143Nd_CD71 |
| 144Nd_CD94 |
| 149Sm_CD127 |
| 155Gd_PD-1 |
| 158Gd_Ki67 |
| 159Tb_CD38 |
| 160Gd_CD14 |
| 164Dy_CD15 |
| 166Er_NKG2D_CD314 |
| 168Er_CD13 |
| 169Tm_NKG2A_CD159 |
| 170Er_pH2AX |
| 172Yb_CD10 |
| 174Yb_CD20 |
| 161Dy_CD16 |
| 146Nd_CD8a |
| 150Nd_CD117 |
| 157Gd_CD45RA |
| 162Dy_CD11b |
| 163Dy_CD64 |
| 171Yb_CD200 |
| 173Yb_CD19 |
| 176Yb_pCREB |
| 115In_CD45 |
| 194Pt_H3K27me3 |
| 147Pm_CD56 |
| 113In_CD3 |
| 209Bi_pHH3 |
| 89Y_CD235 |

## Reference

1. Romain, G. et al. Antibody Fc engineering improves frequency and promotes kinetic boosting of serial killing mediated by NK cells. Blood 124, 3241–3249 (2014).

2. Mani, R. et al. Fc-engineered anti-CD33 monoclonal antibody potentiates cytotoxicity of membrane-bound interleukin-21 expanded natural killer cells in acute myeloid leukemia. Cytotherapy (2020).

3. Gleason, M.K. et al. CD16xCD33 bispecific killer cell engager (BiKE) activates NK cells against primary MDS and MDSC CD33+ targets. Blood 123, 3016–3026 (2014).

4. Vallera, D.A. et al. IL15 Trispecific Killer Engagers (TriKE) Make Natural Killer Cells Specific to CD33+ Targets While Also Inducing Persistence, In Vivo Expansion, and Enhanced Function. Clin Cancer Res 22, 3440-3450 (2016).

5. Dutour, A. et al. In Vitro and In Vivo Antitumor Effect of Anti-CD33 Chimeric Receptor-Expressing EBV-CTL against CD33 Acute Myeloid Leukemia. Advances in hematology 2012, 683065 (2012).

6. Kim, M.Y. et al. Genetic Inactivation of CD33 in Hematopoietic Stem Cells to Enable CAR T Cell Immunotherapy for Acute Myeloid Leukemia. Cell 173, 1439–1453 e1419 (2018).

7. Rotiroti, M.C. et al. Targeting CD33 in Chemoresistant AML Patient-Derived Xenografts by CAR-CIK Cells Modified with an Improved SB Transposon System. Mol Ther 28, 1974–1986 (2020).

8. Liu, E. et al. Use of CAR-Transduced Natural Killer Cells in CD19-Positive Lymphoid Tumors. N Engl J Med 382, 545–553 (2020).

9. Naeimi Kararoudi, M., et al. Genetic and epigenetic modification of human primary NK cells for enhanced antitumor activity. Seminars in Hematology 57, 201–212 (2020).

10. Pomeroy, E.J. et al. A Genetically Engineered Primary Human Natural Killer Cell Platform for Cancer Immunotherapy. Mol Ther 28, 52–63 (2020).

11. Naeimi Kararoudi, M., et al. Generation of Knock-out Primary and Expanded Human NK Cells Using Cas9 Ribonucleoproteins. J Vis Exp (2018).

12. Naeimi Kararoudi, M. et al. CD38 deletion of human primary NK cells eliminates daratumumab-induced fratricide and boosts their effector activity. Blood (2020).

13. Mali, P. et al. RNA-guided human genome engineering via Cas9. Science 339, 823–826 (2013).

14. Suzuki, K. et al. In vivo genome editing via CRISPR/Cas9 mediated homology-independent targeted integration. Nature 540, 144–149 (2016).

15. Liu, J., Zhou, G., Zhang, L. & Zhao, Q. Building Potent Chimeric Antigen Receptor T Cells With CRISPR Genome Editing. Frontiers in immunology 10, 456 (2019).

16. Eyquem, J. et al. Targeting a CAR to the TRAC locus with CRISPR/Cas9 enhances tumour rejection. Nature 543, 113–117 (2017).

17. Moseman, J.E., Foltz, J.A., Sorathia, K., Heipertz, E.L. & Lee, D.A. Evaluation of serum-free media formulations in feeder cell-stimulated expansion of natural killer cells. Cytotherapy 22, 322–328 (2020).

18. Denman, C.J. et al. Membrane-Bound IL-21 Promotes Sustained Ex Vivo Proliferation of Human Natural Killer Cells. PLoS One 7, e30264 (2012).

19. 19. Somanchi, S.S., Senyukov, V.V., Denman, C.J. & Lee, D.A. Expansion, purification, and functional assessment of human peripheral blood NK cells. J Vis Exp (2011).

20. Moseman, J.E., Foltz, J.A., Sorathia, K., Heipertz, E.L. & Lee, D.A. Evaluation of serum-free media formulations in feeder cell-stimulated expansion of natural killer cells. Cytotherapy (2020).

21. Buenrostro, J.D., Giresi, P.G., Zaba, L.C., Chang, H.Y. & Greenleaf, W.J. Transposition of native chromatin for fast and sensitive epigenomic profiling of open chromatin, DNA-binding proteins and nucleosome position. Nat Methods 10, 1213–1218 (2013).

22. Hsiau, T. et al. Inference of CRISPR Edits from Sanger Trace Data. bioRxiv, 251082 (2018).

23. Mendell, J.R. et al. Single-Dose Gene-Replacement Therapy for Spinal Muscular Atrophy. N Engl J Med 377, 1713–1722 (2017).

24. de Vree, P.J. et al. Targeted sequencing by proximity ligation for comprehensive variant detection and local haplotyping. Nat Biotechnol 32, 1019–1025 (2014).

25. Cerignoli, F. et al. In vitro immunotherapy potency assays using real-time cell analysis. PLoS One 13, e0193498 (2018).

26. Behbehani, G.K., Bendall, S.C., Clutter, M.R., Fantl, W.J. & Nolan, G.P. Single-cell mass cytometry adapted to measurements of the cell cycle. Cytometry A 81, 552–566 (2012).

27. Rahman, A.H., Tordesillas, L. & Berin, M.C. Heparin reduces nonspecific eosinophil staining artifacts in mass cytometry experiments. Cytometry A 89, 601–607 (2016).

28. Kotecha, N., Krutzik, P.O. & Irish, J.M. Web-based analysis and publication of flow cytometry experiments. Curr Protoc Cytom Chapter 10, Unit10 17 (2010).

29. Finck, R. et al. Normalization of mass cytometry data with bead standards. Cytometry A 83, 483–494 (2013).

30. Qiu, P. et al. Extracting a cellular hierarchy from high-dimensional cytometry data with SPADE. Nat Biotechnol 29, 886–891 (2011).

31. Schmid-Burgk, J.L., Honing, K., Ebert, T.S. & Hornung, V. CRISPaint allows modular base-specific gene tagging using a ligase-4-dependent mechanism. Nat Commun 7, 12338 (2016).

32. Chen, C.C., Feng, W., Lim, P.X., Kass, E.M. & Jasin, M. Homology-Directed Repair and the Role of BRCA1, BRCA2, and Related Proteins in Genome Integrity and Cancer. Annu Rev Cancer Biol 2, 313–336 (2018).

33. Oceguera-Yanez, F. et al. Engineering the AAVS1 locus for consistent and scalable transgene expression in human iPSCs and their differentiated derivatives. Methods 101, 43–55 (2016).

34. Wang, J. et al. Highly efficient homology-driven genome editing in human T cells by combining zinc-finger nuclease mRNA and AAV6 donor delivery. Nucleic Acids Res 44, e30 (2016).

35. Foust, K.D. et al. Therapeutic AAV9-mediated suppression of mutant SOD1 slows disease progression and extends survival in models of inherited ALS. Mol Ther 21, 2148–2159 (2013).

36. Ran, F.A. et al. Genome engineering using the CRISPR-Cas9 system. Nat Protoc 8, 2281–2308 (2013).

37. MacLeod, D.T. et al. Integration of a CD19 CAR into the TCR Alpha Chain Locus Streamlines Production of Allogeneic Gene-Edited CAR T Cells. Mol Ther 25, 949–961 (2017).

38. He, X. et al. Knock-in of large reporter genes in human cells via CRISPR/Cas9-induced homology-dependent and independent DNA repair. Nucleic Acids Res 44, e85 (2016).

39. Song, F. & Stieger, K. Optimizing the DNA Donor Template for Homology-Directed Repair of Double-Strand Breaks. Mol Ther Nucleic Acids 7, 53–60 (2017).

40. Li, K., Wang, G., Andersen, T., Zhou, P. & Pu, W.T. Optimization of genome engineering approaches with the CRISPR/Cas9 system. PLoS One 9, e105779 (2014).

41. Li, Y., Hermanson, D.L., Moriarity, B.S. & Kaufman, D.S. Human iPSC-Derived Natural Killer Cells Engineered with Chimeric Antigen Receptors Enhance Anti-tumor Activity. Cell Stem Cell 23, 181–192 e185 (2018).

42. McCarty, D.M. Self-complementary AAV vectors; advances and applications. Mol Ther 16, 1648–1656 (2008).

